# The odd one out: *Arabidopsis* reticulon20 has a role in lipid biosynthesis

**DOI:** 10.1101/123679

**Authors:** Verena Kriechbaumer, Lilly Maneta-Peyret, Stanley W Botchway, Jessica Upson, Louise Hughes, Jake Richardson, Maike Kittelmann, Patrick Moreau, Chris Hawes

## Abstract

The family of reticulon proteins has been shown to be involved in a variety of functions in eukaryotic cells including tubulation of the endoplasmic reticulum (ER), formation of cell plates and primary plasmodesmata. Reticulons are integral ER membrane proteins characterised by a reticulon homology domain comprising four transmembrane domains which results in the reticulons sitting in the membrane in a W-topology. Here we report on a subgroup of reticulons with an extended N-terminal domain and in particular on arabidopsis reticulon 20. We show that reticulon 20 is located in a unique punctate pattern on the ER membrane. Its closest homologue reticulon 19 labels the whole ER. We show that mutants in RTN20 or RTN19, respectively, display a significant change in sterol composition in the roots indicating a role in lipid biosynthesis or regulation. A third homologue in this family - 3BETAHSD/D1- is localised to ER exit sites resulting in an intriguing location difference for the three proteins.

## Introduction

The endoplasmic reticulum (ER) is a multifunctional organelle (Hawes *et al.*, 2015) involved in a plethora of aspects of plant life. The polygonal network of the cortical ER consists of motile tubules that are capable of morphing into small cisternae, mainly at the three-way junctions of the ER network (Sparkes *et al.*, 2009). The plant cortical ER network has been shown to play numerous roles in protein trafficking (Palade, 1975; Vitale and Denecke, 1999) and pathogen responses (Pattison and Amtmann, 2009; Beck *et al.*, 2012). It is a highly dynamic organelle and previous studies have demonstrated a possible link between ER structure and function within different cell- and tissue-types (Stephenson and Hawes, 1986; Lai *et al.*, 2014).

A variety of ER movements have been characterised, including growth and shrinkage of tubules, rearrangement of the polygonal network (Sparkes *et al.*, 2009), movement of the membrane surface (Runions *et al.*, 2006), and the conversion between cisternal and tubular ER (Hawes *et al.*, 2015). These distinct movements which appear dependent on the acto/myosin system (Sparkes *et al.* 2009) and the possibly significant link between structure and function, makes these processes important to understand.

### The reticulon protein family

Reticulons are a family of proteins found in a wide range of eukaryotes and have been shown to localise to the ER in many species, including mammals, yeasts and plants (Nziengui *et al.*, 2007; Yang and Strittmatter, 2007). Previous studies have demonstrated a role for these proteins in moving and shaping the ER into tubules (Voeltz *et al.*, 2006; Yang and Strittmatter, 2007; Tolley *et al.*, 2008).

Plant reticulons (RTNLB-reticulon-like protein B; henceforth referred to as RTN) are considered to be essential in maintaining the tubular ER network as they contribute significantly to tubulation of the ER (Tolley *et al.*, 2008, 2010; Chen *et al.*, 2012). In arabidopsis, the reticulon protein family comprises 21 members (Oertle *et al.*, 2003; Nziengui *et al.*, 2007). Despite overlapping functions of the members of this protein family, variation in reticulon isoform expression does occur within different tissues. For example, AtRTN13 was found to be more abundant in the seeds compared with the rest of the plant (Sparkes *et al.*, 2010), suggesting there may be cell-specific roles for RTN isoforms (Yang and Strittmatter, 2007). RTNs are integral ER membrane proteins characterised by a C-terminal reticulon homology domain (RHD) which has been suggested to generate and/or stabilize curvature of the membrane. This conserved domain of about 200 amino acids contains two long hydrophobic regions flanking a hydrophilic loop. The hydrophobic regions can each be further subdivided into two transmembrane domains (TMDs) resulting in a ‘W’–like topology. The N-and C-termini of the protein are facing the cytosol (Sparkes *et al.*, 2010). Reticulon proteins can dimerize or oligomerize and thereby cause localized tensions in the ER membrane inducing membrane curvature (Sparkes *et al.*, 2010). When overexpressed in planta, RTNs induce severe constrictions of ER tubules and are able to convert ER membrane sheets into tubules (Tolley *et al.*, 2008, 2010: Sparkes *et al.*, 2010). More recently AtRTN13 was shown to harbour an amphipathic helix at its cytosolic C-terminus which also appears to be involved in inducing ER membrane curvature (Breeze *et al.*, 2016).

This membrane constriction of RTNs has also been shown to be of importance in the context of cell plate formation and primary plasmodesmata (Knox *et al.*, 2015). Plasmodesmata formation is dependent on tubulating the cortical ER to form the desmotubules, axial structures of 15 nm diameter crossing the plasmodesmata pore and thereby connecting the ER of two cells (Overall and Blackman, 1996; Ehlers and Kollmann, 2001; Tilsner *et al.*, 2011). Two reticulon proteins (RTN3 and RTN6) were identified in a proteomic study as plasmodesmata-localised proteins (Fernandez-Calvino *et al.*, 2011) and could also be shown recently to be present in primary plasmodesmata at cytokinesis (Knox *et al.*, 2015). RTN3 and 6 also interact specifically with themselves and each other and a variety of plasmodesmata proteins and proteins previously described as targets of viral movement proteins (Kriechbaumer *et al.*, 2015).

A third reticulon protein predicted to be plasmodesmata localised is RTN20 (TAIR, https://www.arabidopsis.org). RTN20 is one of 5 reticulon proteins (RTN17-21) that features an additional N-terminal domain predicted to have an enzymatic function in sterol biosynthesis (AraCyc).

### Plant sterols

Sterols are essential in all eukaryotic cellular membranes but their biosynthesis differs greatly between animals, plants and fungi. Plant sterols such as sitosterol, stigmasterol and campesterol influence the permeability and fluidity of membranes by lipid-lipid as well as lipid-proteins interactions within the membrane (Hartmann 1998). Enhanced sterol levels can often be detected in detergent-insoluble membrane rafts in the plasma membrane (Borner *et al.*, 2005; Cacas *et al.*, 2012) which are suggested to be important for signalling processes (Mongrand *et al.*, 2010; Simon-Plas *et al.*, 2011) involving for example auxin transport (Titapiwatanakun *et al.*, 2009) and PAMPs (pathogen-associated molecular patterns, Bhat *et al.*, 2005). Sterol molecules become functional after removal of the two methyl groups at the C-4 position of cycloartenol, a precursor molecule of plant sterols. The enzymatic activities required for this C-4 demethylation in plants have been characterized: this requires the activity of a sterol C-4 methyl oxidase (Pascal *et al.*, 1993; Darnet and Rahier, 2004), a 3beta-hydroxysteroid dehydrogenase/C-4 decarboxylase (3BETAHSD/D) (Rondet *et al.*, 1999) and an NADH-dependent 3-oxosteroid reductase (Pascal *et al.*, 1994). One of the recently characterized 3BETAHSD/D2 proteins (Rahier *et al.*, 2006; 3BETAHSD/D2) is actually a reticulon protein (RTN19). By using a three-dimensional homology modeling to identify key amino acids, it has been determined that this protein is a bifunctional short-chain dehydrogenase/reductase enzyme (Rahier et al, 2009; Rahier et al, 2013).

Here we show a unique localisation for RTN20 on the ER membrane as well as an indication for a role in lipid biosynthesis or regulation. We show that mutants in RTN20 or RTN19, respectively, display a significant change in sterol composition in the roots and that a third homologue in this family - 3BETAHSD/D1- is localised to ER exit sites resulting in an intriguing location difference for the three homologues.

## Results

### Reticulon phylogeny and sub-classes

The reticulon protein family in *Arabidopsis thaliana* consists of 21 family members. The proteins group according to structural organisation of the functional domains, with those proteins mainly consisting of the reticulon homology domain (RTN1-16) grouping together but clearly differentiated from the reticulons with an additional N-terminal domain (RTN17-21) (Fig. 1). Within the group of reticulons containing the additional N-terminal domain RTN19 and 20 are again phylogenetically differentiated from RTN17, 18, and 21.

**Figure 1:**
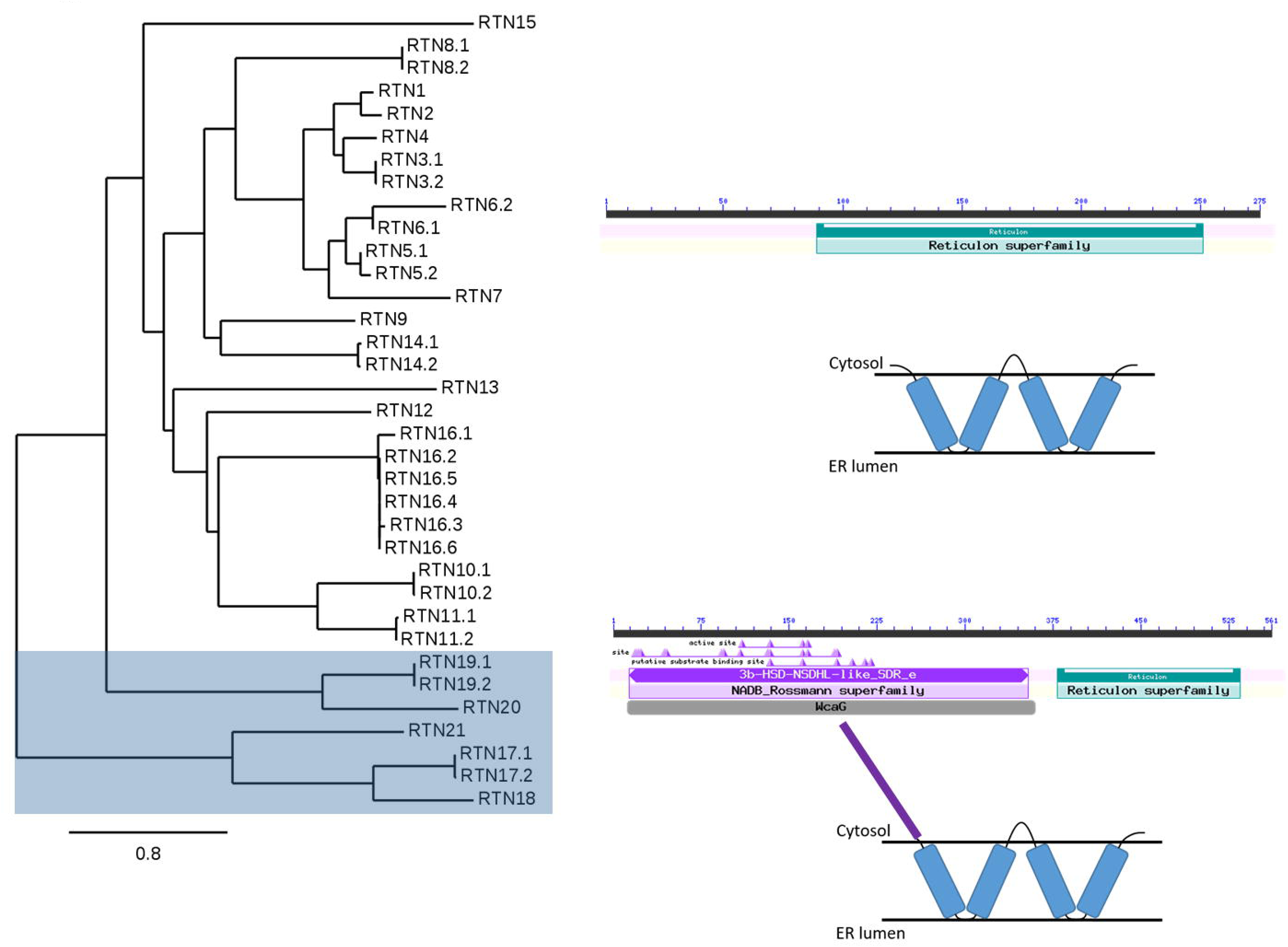
Phylogenetic analysis of reticulon proteins. Phylogenetic relationships within the arabidopsis reticulon family including alternate splice isoforms are shown. Reticulon proteins with an extended N-terminal domain (RTN17-21) are highlighted with a grey shade (left). Bar for bootstrap value is shown. BLAST domain annotations and membrane topology diagrams for this group as well as for the other reticulon family proteins (RTN1-16) are indicated (right).

### RTN20 subcellular location

RTN20 tagged to the yellow fluorescent protein was transiently expressed in tobacco leaf epidermal cells using Agrobacterium-mediated transformation (Fig. 2 A). YFP-RTN20 labels the ER but rather than showing the typical ER tubule phenotype, the expression pattern appears as dots along the ER (Fig. 2A). The same unusual expression pattern could be observed when RTN20 was stably transformed into *Arabidopsis thaliana* (Fig. 2 B, C).

**Figure 2:**
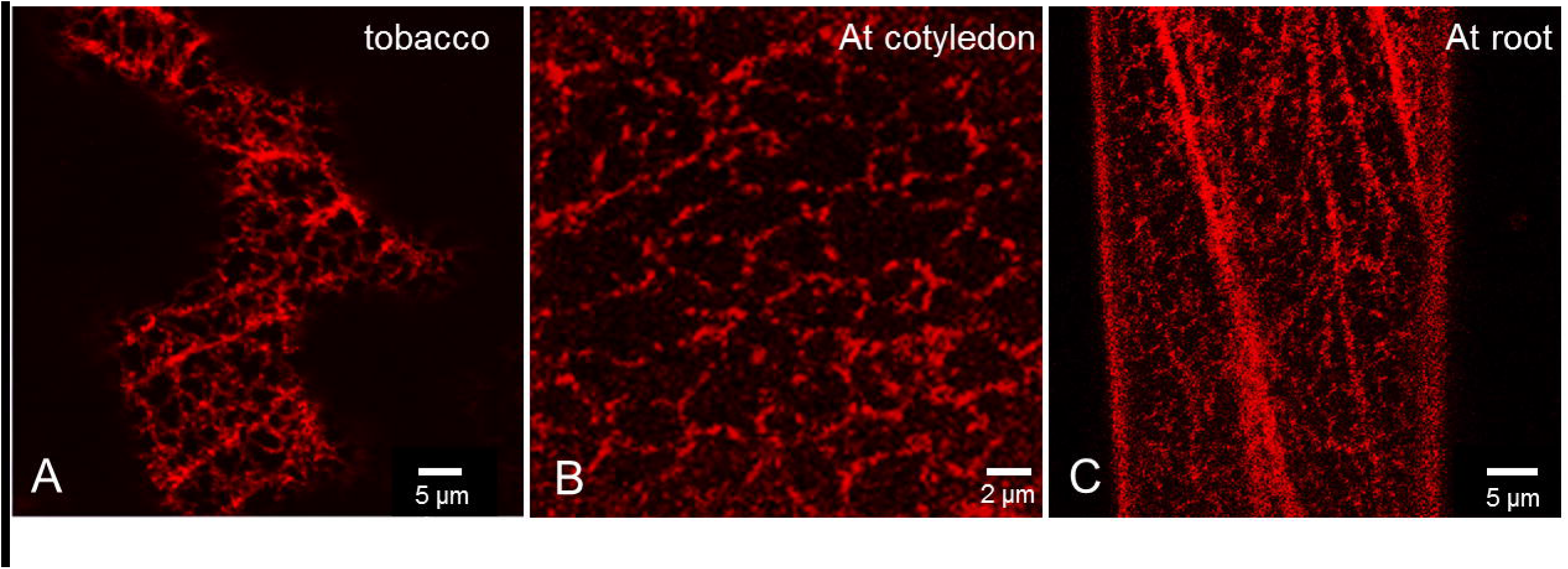
Comparison of RTN20 expression in tobacco and arabidopsis. Expression of YFP-RTN20 is shown transient in tobacco leaf epidermal cells (A) and in a stable manner in Arabidopsis thaliana cotyledon cells (B) and root cells (C).

High resolution Airyscan imaging of GFP-RTN20 together with RFP-RTN1 clearly shows the dotted structure of RTN20 on the ER membrane (Fig. 3 A). RTN20 also does not exhibit the typical reticulon constriction phenotype previously reported (Sparkes *et al.*, 2010; Breeze *et al.*, 2016) when coexpressed with GFP-HDEL (Fig. 3 B)

**Figure 3:**
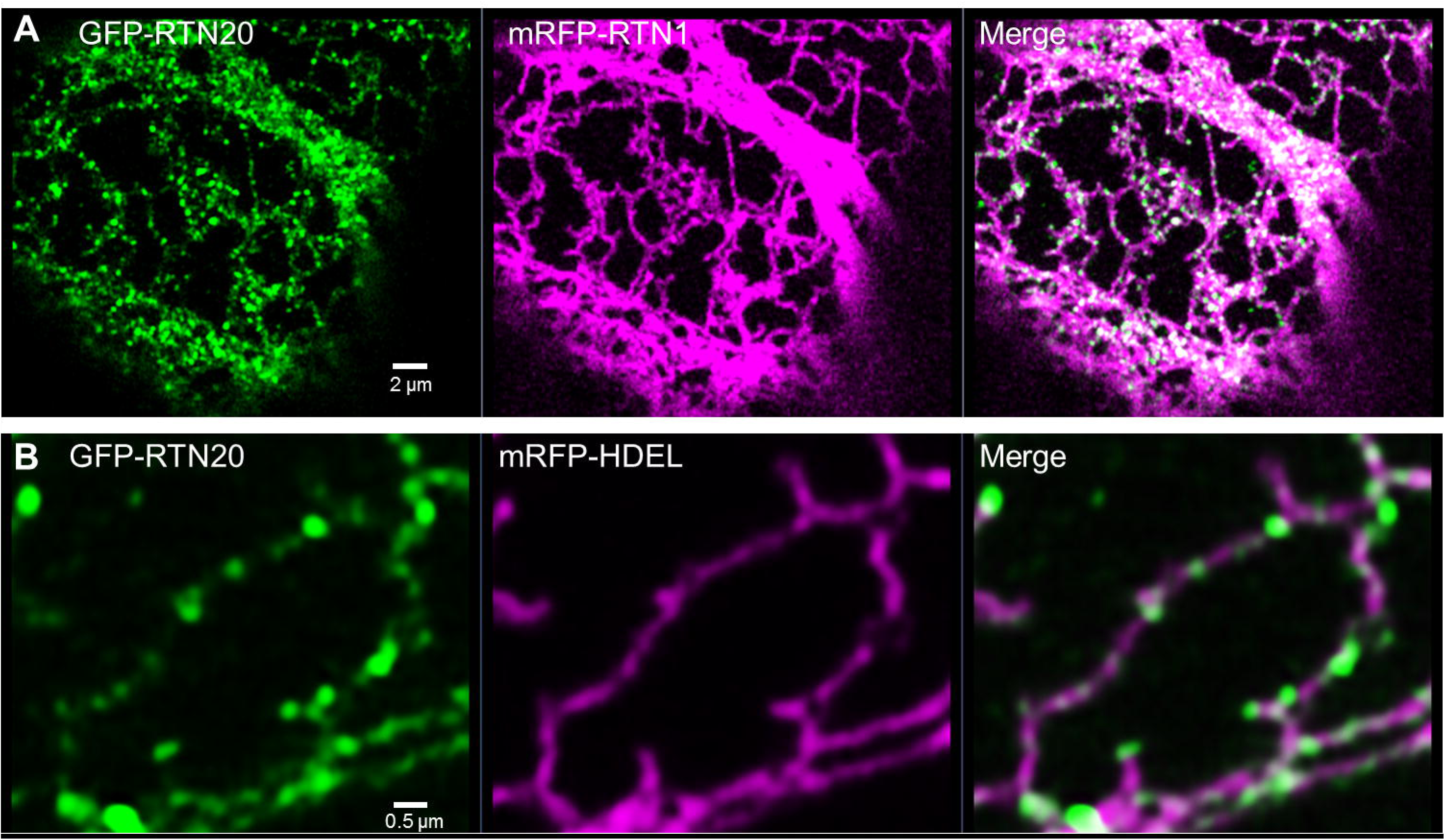
Airyscan confocal images for RTN20 subcellular localisation. GFP-RTN20 is expressed in tobacco leaf epidermal cells by Agrobacterium-mediated transient expression together with RFP-RTN1 (A) or RFP-HDEL (B), respectively. Size bars are given.

To determine if the localisation of RTN20 is a feature for the group of reticulons with extended N-terminal domain, subcellular localisation of RTN19 was also tested as this is the most closely related reticulon to RTN20. RTN20 and RTN19 share 42% amino acid identity. In confocal imaging RTN19 clearly labels the ER when co-expressed with either RTN20 or GFP-HDEL, respectively, but shows classic ER membrane labelling as other reticulons (Sparkes *et al.*, 2008) rather than the punctate expression pattern along the ER of RTN20 (Fig. 4).

**Figure 4:**
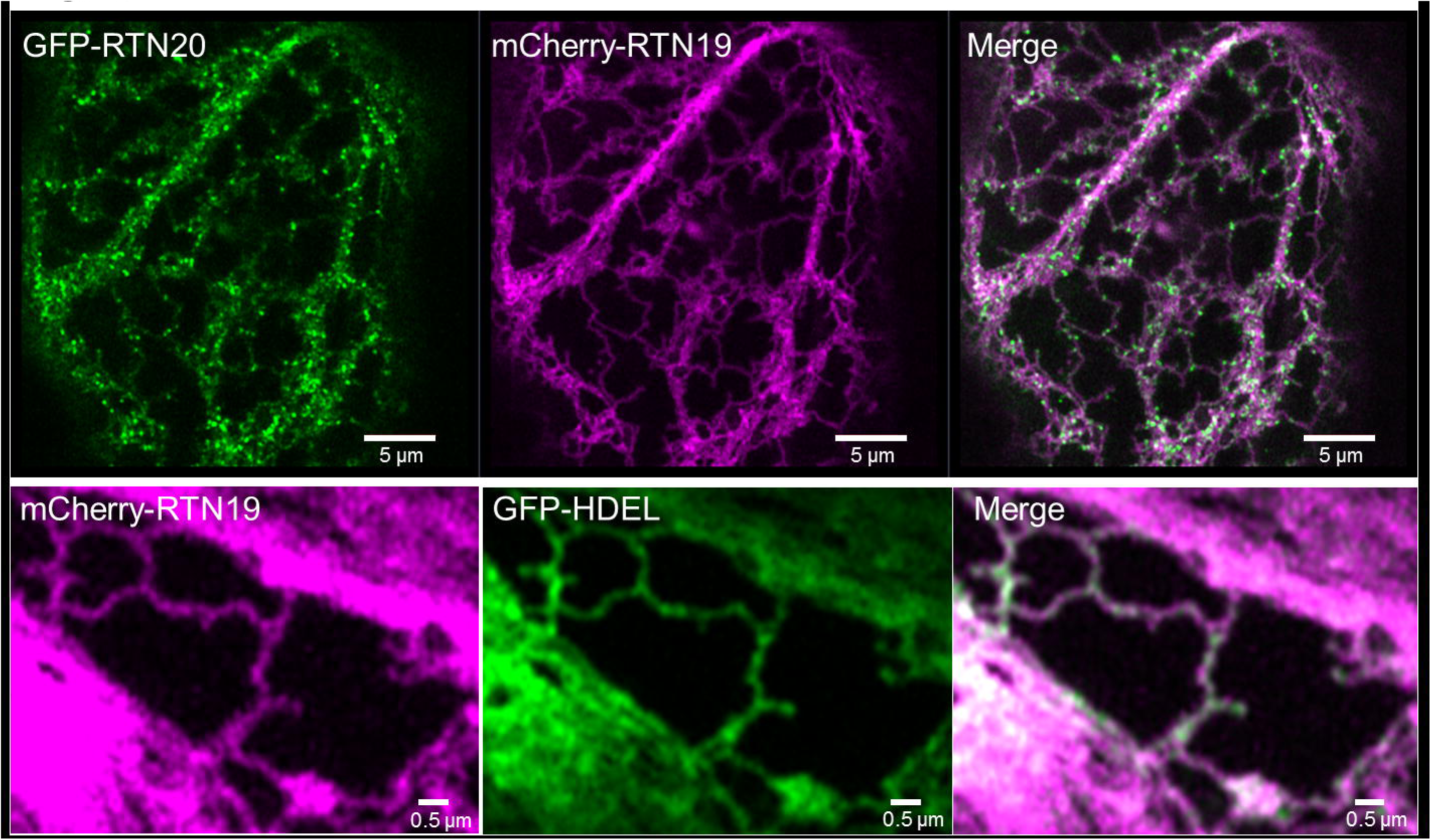
Confocal images for RTN19 localisation. RTN19 fused to mCherry is coexpressed in tobacco leaf epidermal cells with RTN20 (top row) as well as the ER-marker GFP-HDEL (bottom row). RTN19 shows colocalisation with HDEL but not the dotted pattern of RTN20. Size bars are given.

To test what could cause this difference in localisation between RTN20 and RTN19 on the amino acid level fusions between the N-terminus of RTN19 and the C-terminus of RTN20 were created (Supplementary Figure S1). Fusing the RTN20 C-terminus including the transmembrane domain area as well as the C-terminal tail to the N-terminus of RTN19 (lacking its TMDs) resulted in RTN19 localising to punctate structures similar to RTN20 localisation (Supplementary Figure S1 A). The same change in localisation happened when only the C-terminal tail without the TMDs from RTN20 was fused to the N-terminus of RTN19 with its TMD (Supplementary Figure S1 B).

In order to assess whether the punctate distribution of YFP-RTN20 had any effect on ER morphology, we carried out an electron microscope analysis of ER structure in YFP-RTN20 expressing arabidopsis plants and in wild type plants. As leaf cells are very large and therefore difficult to use for EM analysis, we used root tip cells as our experimental material. To visualise the ER network, ER was selectively stained using the zinc iodide osmium tetroxide impregnation technique and reconstructed in 3-D by serial block face scanning electron microscopy (Kittelmann *et al.*, 2016). No major differences were observed in ER structure between wild type and YFP-RTN20 expressing plants (Supplementary Figure S2) suggesting that the punctate distribution of fluorescence most likely reflects clustering of the protein in patches on the ER membrane. Also despite the root lipid phenotype in the *rtn20* mutant plants no significant differences could be observed in mutant roots on EM level (Supplementary Figure S2).

### Protein interactions between RTN 20 and other reticulon proteins

It has been shown that reticulon proteins are capable of forming homomers as well as dimers with other reticulon proteins (Sparkes *et al.*, 2010). Due to the difference in the N-terminal domain structure, it was of interest to test if RTN20 is still capable of such interactions. Förster resonance energy transfer by fluorescence lifetime imaging microscopy (FRET-FLIM) analysis to confirm prey-bait interactions *in vivo* was applied (Kriechbaumer *et al.*, 2015). This technique enables the measurement of the space map of picosecond fluorescence decay at each pixel of the image through confocal single and multiphoton excitation. Förster resonance energy transfer (FRET) used to determine the colocalisation of two colour chromophores is used to define physical interactions of protein pairs tagged with appropriate GFP fluorophores and monomeric red fluorescent protein (mRFP).

In case of a positive protein-protein interactions FRET-FLIM shows reductions in the excited-state lifetime of GFP (donor) fluorescence in the presence of the acceptor fluorophore (mRFP). A reduction in fluorescence lifetime of the donor indicates that the two tested proteins are within a distance of 10 nm or less and thereby further indicating a physical interaction between the two proteins (Sparkes *et al.*, 2010; Schoberer and Botchway, 2014; Kriechbaumer *et al.*, 2015).

Due to limitations in the speed of photon counting of the FLIM set-up, measurements are best to be taken from high-expressing regions of ER showing relatively low mobility, such as the ER associated with the nuclear envelope. This allows more reliable measurements than the fast-moving cortical ER (Sparkes *et al.*, 2010) and the use of actin depolymerising agents such as Latrunculin B which can perturb ER structure can be avoided.

For this GFP-RTN20 was used as a donor and the reticulon proteins RTN1, 2, 3, and 19 fused to mRFP as acceptor proteins. These reticulon proteins were chosen to represent the various sub-groups of reticulons known to date: RTN1, 2, and 3 feature the typical reticulon domain topology whereas RTN19 features an additional N-terminal enzymatic domain similar to RTN20. Furthermore RTN3 –together with RTN6- has been shown to be localised to plasmodesmata (Knox *et al.*, 2015). For each combination at least two biological samples with a minimum of three technical replicates each were used for the statistical analysis. FRET-FLIM interaction results are shown in Table 1, Figure 5 and Supplementary Figure S3. Live cells expressing RTN20 alone without an acceptor present was used as a control and resulted in a baseline fluorescence life time of 2.4 ± 0.03 ns. Excited-state lifetimes determined for all RTN-RTN heteromeric interactions tested showed an average fluorescence lifetime of 2.2 ns (Table 1, Fig. 5, Supplementary Figure S3), which is 0.2 ns lower than the donor alone and statistically significantly different from that of the GFP alone.

**Table 1.** Fluorescence lifetimes in FRET-FLIM analysis. Donor and acceptor protein constructs are indicated together with the average fluorescence lifetime (in ns) for the donor fluorophore and the Standard Error (SE) for each combination. The reduction in fluorescence lifetime of the donor is indicated (Δ). It was shown previously that a reduction in excited-state lifetime of 0.2 ns is indicative of energy transfer (Stubbs *et al.*, 2005). For each combination, at least two biological samples with a minimum of three technical replicates were used for the statistical analysis.

| **Donor (GFP)** | **Acceptor (mRFP)** | **Average [ns]** | **SE** | **$\Delta$ [ns]** |
| --- | --- | --- | --- | --- |
| RTN20 | (-) | 2.4 | 0.03 | 0.0 |
|  | RTN1 | 2.2 | 0.02 | 0.2 |
|  | RTN2 | 2.2 | 0.02 | 0.2 |
|  | RTN3 | 2.2 | 0.04 | 0.2 |
|  | RTN19 | 2.2 | 0.06 | 0.2 |

**Figure 5:**
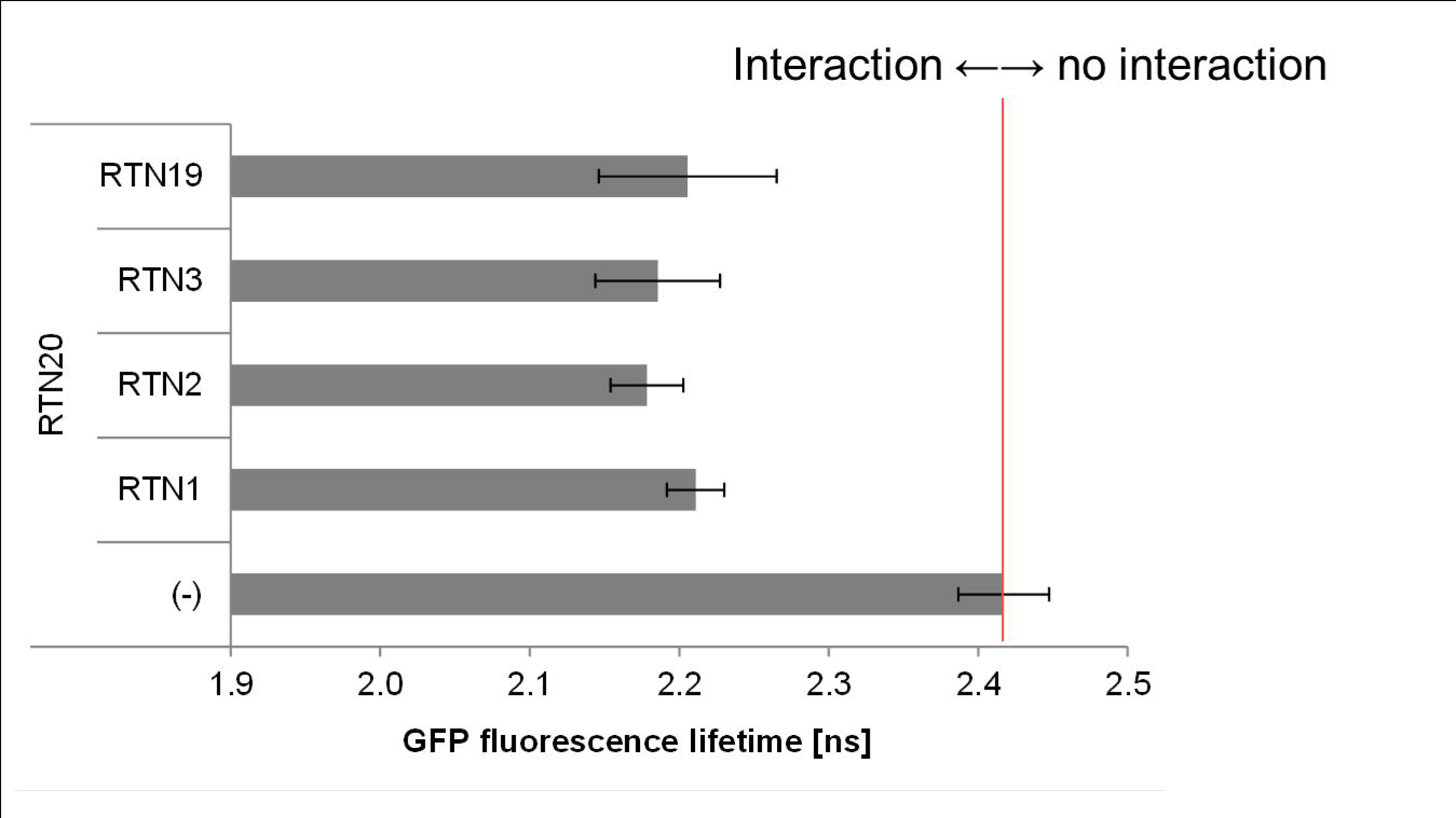
RTN20 protein interactions measured by FRET-FLIM. Bar chart of fluorescent lifetimes for the donor GFP-RTN20 alone as a negative control and in heteromerisation with RTN1, 2, 3, 19 fused to mRFP as acceptor.

Figure 6 shows the FRET-FLIM analysis steps for GFP-RTN20 alone (Fig. 6 A-D) as a control and for GFP-RTN20 interacting with mRFP-RTN1 (Fig. 6 E-I) as an example for interaction. Raw FRET-FLIM images are shown in Figure 6 A and E. This analysis takes into account the lifetime values of each pixel within the image visualized by a pseudocoloured lifetime map (Fig. 6 B and F). The graph shows the distribution of lifetimes within the image (Fig. 6 C and G), with blue shades representing longer GFP fluorescence lifetimes than green/yellow ones. Decay curves (Fig. 6 D and H) of representative single pixel highlight an optimal single exponential fit, where X^2^ values in the range of 0.9 to 1.2 were considered an excellent fit to the data. Confocal images for the region of interest showing the GFP construct in green and the mRFP construct in red are shown. This specific example shows that RTN20 heterodimerizes with RTN1 as the lifetime values for the GFP/mRFP fusion pair (2.21 ± 0.02 ns; Table 1) are significantly lower than those for the GFP fusion alone (2.42 ± 0.03 ns).

**Figure 6:**
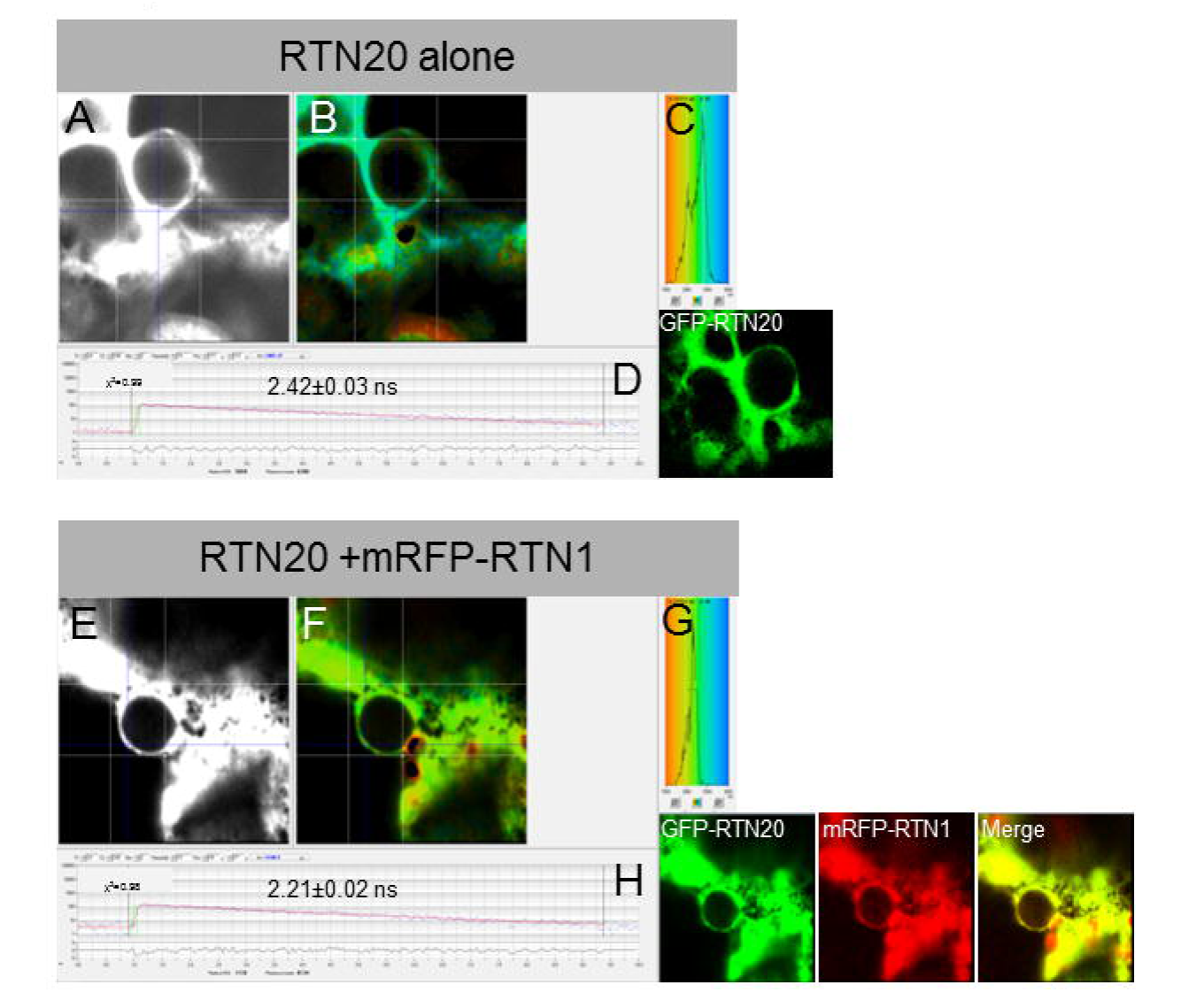
FRET-FLIM analysis of RTN20. FRET-FLIM analysis of RTN20 without an interaction partner (A–D) or RTN20-RTN1 dimerization (E–I) is shown. A and E show the raw FRET-FLIM data. In B and F, pseudocoloured lifetime maps display the lifetime values for each point within the region of interest. The distribution of lifetimes across the entire image is shown in C and G with blue shades representing longer GFP fluorescence lifetimes than green ones. D and H show representative decay curves of a single point with an optimal single exponential fit, where X^2^ values from 0.9 to 1.2 are considered an excellent fit to the data points. Respective confocal images for the analysis are given with the GFP construct in green and the mRFP construct in red. This example of FRET-FLIM analysis shows that RTN20interacts with RTN1 because the lifetime values for the GFP/mRFP fusion pair (H; 2.21 ± 0.02 ns) are lower than those for the GFP fusion alone (D; 2.42± 0.03 ns).

### Lipid analysis in rtn20 mutants

As the N-terminus of RTN20 is predicted to be involved in sterol biosynthesis, lipid composition in the *rtn20* mutant was tested. The closest homologue to RTN20 - RTN19- has previously been shown to possess 3beta-hydroxysteroid dehydrogenase/C-4 decarboxylase (3BETAHSD/D) activity (Rahier *et al.*, 2006) and the *rtn19* mutant was therefore included in the lipid analysis. Furthermore a mutant rescue line was created by overexpressing RTN20 under a 35S promoter in the *rtn20* mutant background

Sterols and the major phospholipids phosphatidylcholine (PC) and phosphatidylethanolamine (PE) from roots and leaves of two week-old seedlings (wild type, mutant lines *rtn20* and *rtn19*, and rescue line) were determined from 3 independent experiments by HPTLC coupled to densitometry. The amounts of sterols, PC and PE in the roots and leaves of control, mutant and rescue lines were determined as μg/g FW, and the values for the control roots and leaves were taken as equal to 100 and the corresponding values for the mutant and rescue lines were calculated accordingly (Fig. 7 A). The ratios Sterols to PC+PE are also indicated for the different lines (Fig. 7 B).

**Figure 7:**
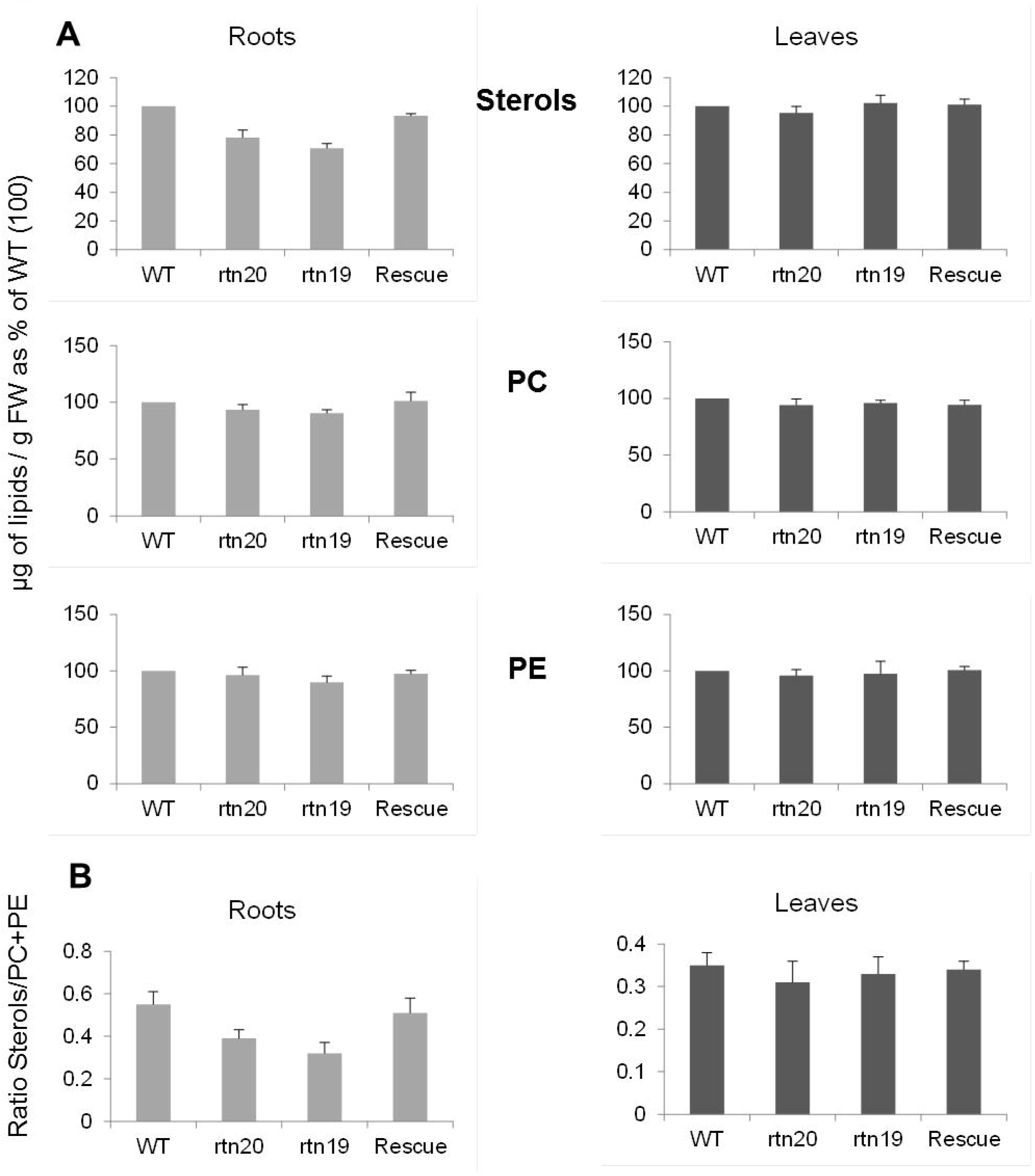
Lipid composition of the reticulon mutants *rtn19* and *rtn20* and rescue lines. (A) Sterols and the major phospholipids phosphatidylcholine (PC) and phosphatidylethanolamine (PE) from roots and leaves of two week-old seedlings (wild type, mutant lines *rtn20* and *rtn19*, and rescue line) were quantified. Data was obtained from 3 independent experiments by HPTLC coupled to densitometry. Sterol, PC and PE amounts were determined as μg/g FW, the values for the control roots and leaves were normalized to 100 and the corresponding values for the mutant and rescue lines were calculated accordingly. (B) The ratios between sterols to PC+PE content are indicated for the different lines.

The levels of sterols and that of the major phospholipids phosphatidylcholine (PC) and phosphatidylethanolamine (PE) from roots and leaves of two week-old seedlings (wild type and mutant lines) were determined from three independent experiments by HPTLC coupled to densitometry as indicated in the experimental section. We determined the following amounts of sterols, PC and PE in the control roots (461 ± μg/g FW of sterols; 591 ± 74 μg/g FW of PC and 762 ± 81 μg/g FW of PE) and control leaves (357 ± 23 μg/g FW of sterols; 661 ± 86 μg/g FW of PC and 547 ± 48 μg/g FW of PE). As presented in Figure 7 A, the values for the control roots and leaves were taken as equal to 100 and the corresponding values for the *rtn19*, the *rtn20* and the rescue lines were calculated accordingly. As indicated in Figure 7 A, a significant decrease of the amounts of sterols was observed in the roots of the *rtn19* and *rtn20* mutants (p<0.01) but no variation of the sterols was observed in the leaves of the *rtn19* and *rtn20* mutants, and no significant variation of the phospholipids PC and PE was observed both in the roots and the leaves of the *rtn19* and *rtn20* mutants. This led to a decrease of the sterols to phospholipid (PC+PE) ratio in the roots of the *rtn19* and *rtn20* mutants (Fig. 7 B; p<0.01). Expressing the *RTN20* protein in the mutant background restored the level of sterols (Fig. 7 A; p>0.2) and the sterols to phospholipids (PC+PE) ratio in the roots of the rescue lines (Fig. 7B; p>0.5). Therefore, our results highly suggest that *RTN19* and *RTN20* have an impact on the sterols content of arabidopsis roots.

As the mutant lipid testing showed different effects between roots and leaves *RTN20* transcript levels were examined. To test for the presence of *RTN20* mRNA reverse transcriptase PCR was performed (Fig. 8) using root and cotyledon tissue from both wild-type (WT) Col0 plants and *rtn20* mutant seedlings. *RTN20* could be detected in wildtype root tissue but not in cotyledon tissue (Fig. 8, lane 1 and 2) which could account for the change in lipid composition in *rtn20* roots but not in cotyledon tissue.*rtn20* mutant plants show no detectable *RTN20* transcript (Fig. 8, lane 3 and 4). To test for the quality of the cDNA *rtn20* mutants were also probed with primers for *RTN6* resulting in a *RTN6* band (Fig. 8, lane 5) that was not detectable in the *rtn6* mutant background (Fig. 8, lane 6).

**Figure 8:**
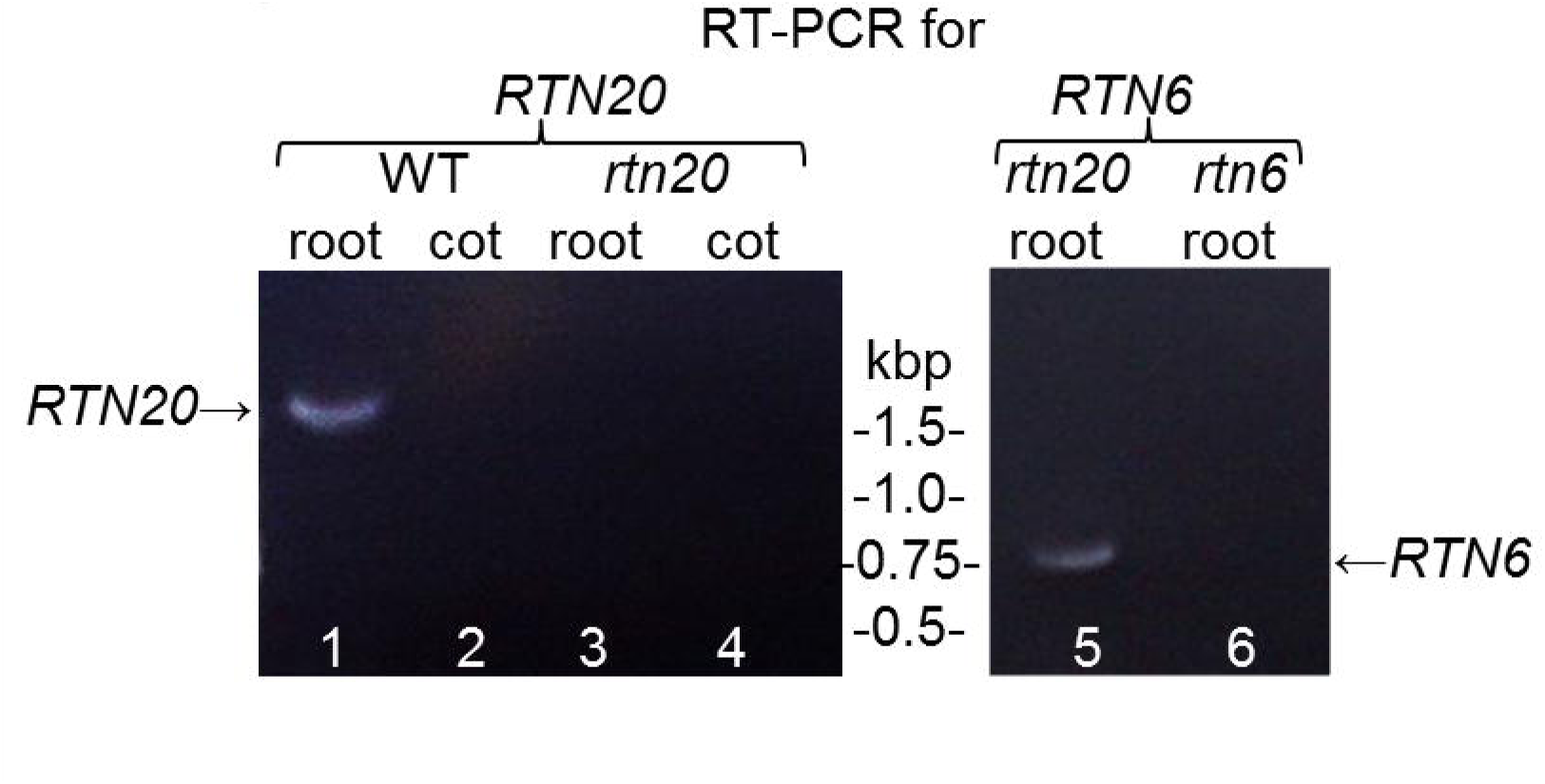
Transcription of RTN20. Reverse transcriptase PCR was performed in wild-type (WT) Col0 plants as well as *rtn20* and *rtn6* mutant plants. It was also distinguished between root and cotyledon (cot) tissue. *RTN20*-mRNA was detectable in WT roots but not in cotyledon tissue or in the *rtn20* mutant. *RTN6* could be detected in the *rtn20* mutant but not in the *rtn6* mutant.

### “The third man”: 3BETAHSD/D1, a homologue of RTN20 and RTN19

Phylogenetic analysis revealed that arabidopsis RTN20 is highly homologous to the yeast protein Erg26p and the mammalian protein sterol-4-α carboxylate 3- dehydrogenase (Fig. 9); both of these analogous proteins have a role in sterol biosynthesis (Gachotte *et al.*, 1998; Caldas and Herman, 2003). Additionally, these proteins were found to be homologous to the arabidopsis 4-α carboxysterol-C3-dehydrogenase (3beta-hydroxysteroid-dehydrogenase/decarboxylase isoform 1,3BETAHSD/D1; Rahier *et al.*, 2006), also involved in sterol biosynthesis. Phylogenetic analysis demonstrated that the newly identified proteins all grouped with RTN19 and RTN20 rather than with the other reticulon proteins with an additional N-terminal domain (RTN17, 18, 21) (Fig. 9), indicating these reticulons may have similar functions. Unlike for the reticulon proteins 19 and 20, no transmembrane domains were predicted for 3BETAHSD/D1 (Supplementary Figure S4).

**Figure 9:**
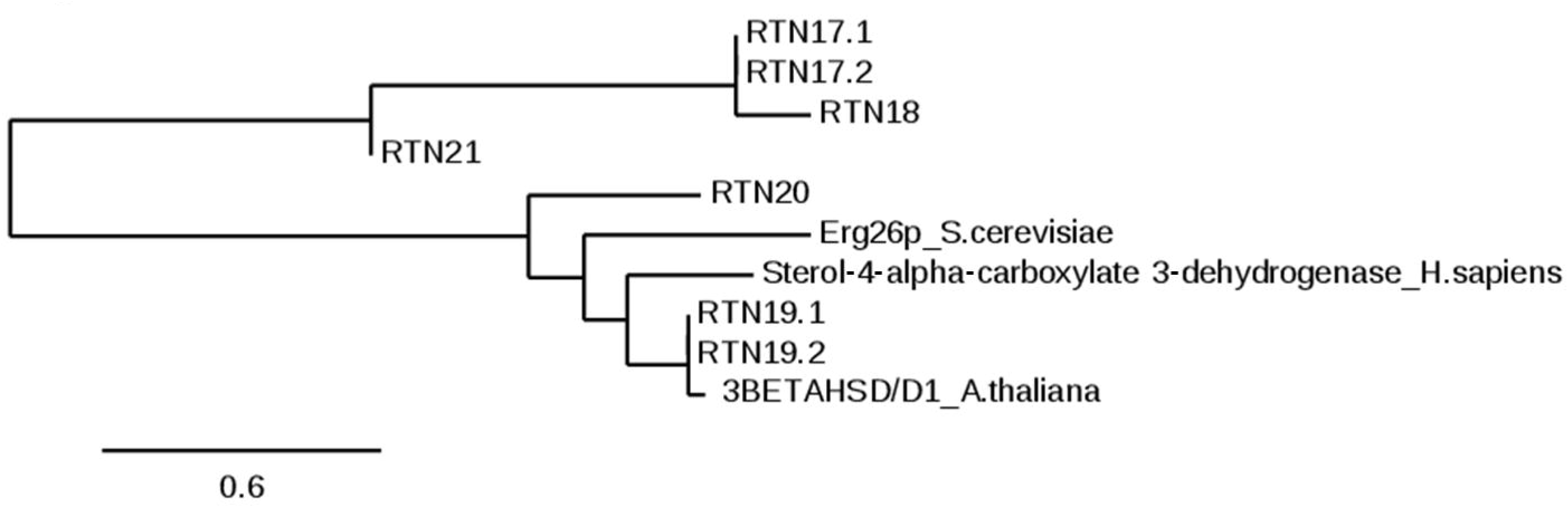
Phylogenetic analysis of RTN20 homologue proteins. A phylogenetic analysis with reticulon proteins featuring an additional N-terminal domain (RTN17-21) together with the closest mammalian homologues is shown. Bar for bootstrap value is shown.

Over the aligned sequence part 3BETAHSD/D1 shows 46% amino acid identity with RTN20 and 82% identity with RTN19 (BLASTP).

3BETAHSD/D1 is not being predicted to have any hydrophobic transmembrane domain or signal peptide but features a potential ER retrieval signal at the C-terminus (KKID, Rahier *et al.*, 2006). Transformation of the yeast ergosterol *erg26* mutant, which lacks 3BETAHSD/D activity, with the arabidopsis 3BETAHSD/D1 gene can complement the mutation and interestingly the activity can be found in microsomal extracts; cytosolic fractions failed to show activity, leading to the speculation that the enzyme is membrane-bound (Rahier *et al.*, 2006). Disruption of ERG26 is lethal, and the *erg26* strain requires ergosterol or cholesterol supplementation for viability pointing to a role in sterol biosynthesis for 3BETAHSD/D1 (Gachotte *et al.*, 1998). Neither single nor double knockout plants of *RTN19* and/or 3BETAHSD/D1 display a visible phenotype (Kim *et al.*, 2012).

When expressing 3BETAHSD/D1 fused to the red fluorescent protein mCherry transiently in tobacco epidermal leaf cells the fusion protein showed neither cytosolic or ER localisation but distinct punctae (Fig. 10). To determine the subcellular localisation of 3BETSHSD/D1 the protein was coexpressed with markers labelling the Golgi bodies or ER exit sites, respectively (Fig. 10). Imaging with a high resolution Airyscan detector it can be seen that 3BETAHSD/D1 only partially colocalises with the *trans*-Golgi marker ST-GFP (Fig. 10 A) and displays more of a doughnut pattern with the Golgi construct in the middle. 3BETSHSD/D1 colocalises better with ER exit site markers Sec16 (Fig. 10 B) and AtSAR1A (Fig. 10 C) all constructs labelling punctate ring-like structures. 3BETSHSD/D1 does not colocalise with the RTN20 punctuate structures (Supplementary Figure S5).

**Figure 10:**
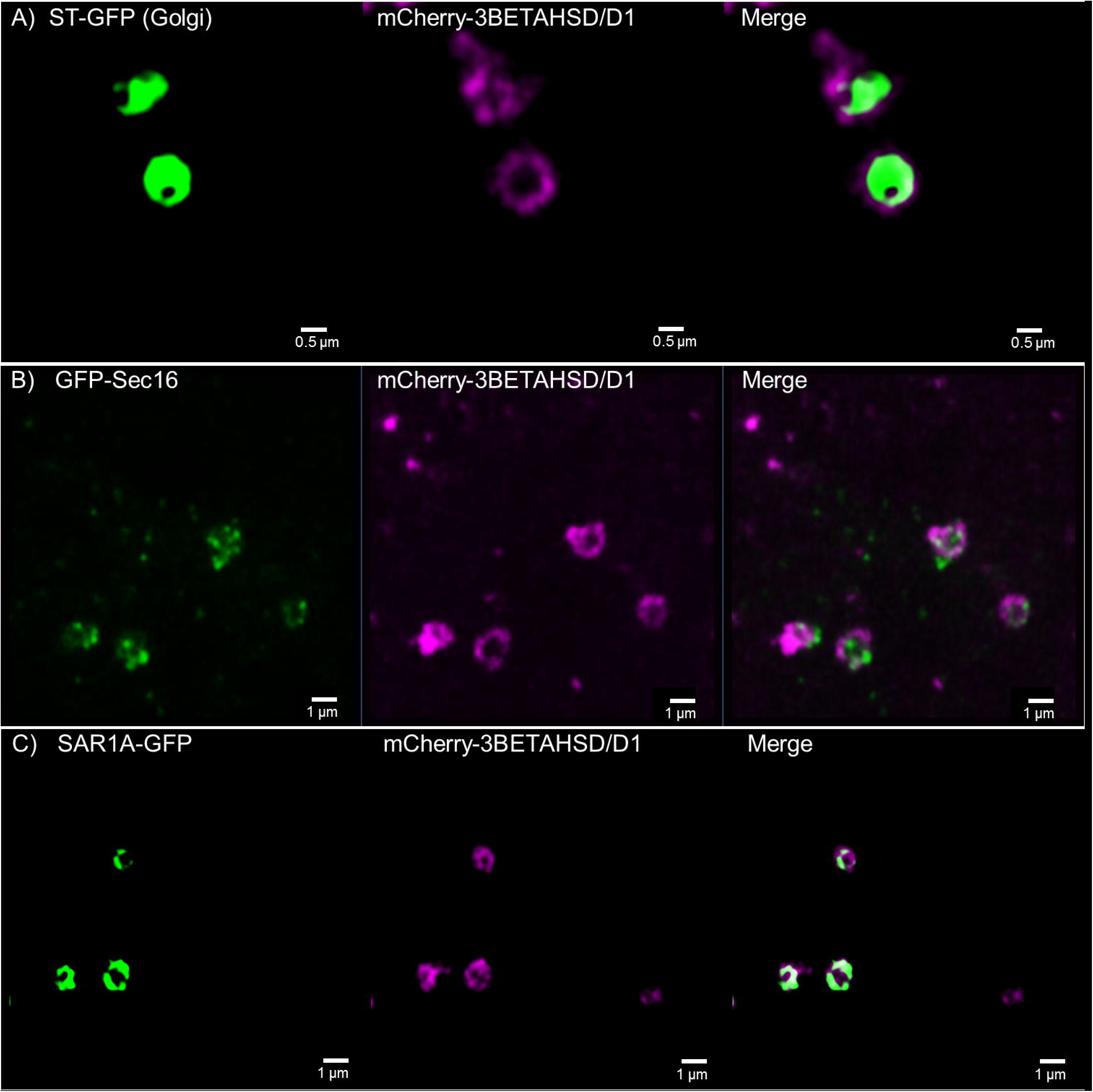
Confocal images for 3BETAHSD/D1 subcellular localisation. 3BETAHSD/D1 fused to the mCherry-fluorescent protein is coexpressed in tobacco leaf epidermal cells with the Golgi marker ST-GFP (A) as well as the ER exit site markers GFP-Sec16 (B) and SAR1A-GFP (C). 3BETAHSD/D1 shows co-localisation with both ER exit site markers but resembles with the ring-like structure more the SAR1A pattern than the dottier Sec16. 3BETAHSD/D1 also partially colocalises with the Golgi marker but circles the marker. Size bars are given.

## Discussion

### Phylogeny and conserved domains in the reticulon protein family

Within the 21 reticulon family members in arabidopsis five proteins cluster together due to an additional N-terminal domain from the reticulon homology domain that all reticulon proteins feature (Fig. 1).The significance of the reticulon homology domain and the C-terminal domain for the ability of reticulon proteins to induce membrane curvature has been demonstrated (Sparkes *et al.*, 2010; Breeze *et al.*, 2016). The reticulon homology domain has been shown to be required for reticulon residence in high-curvature ER membranes and ER tubule constriction, yet it is not necessary for reticulon oligomerisation (Sparkes *et al.*, 2010). A predicted amphipathic helix (APH) at the C-terminus of *RTN13* has been shown to be responsible for tubule constriction (Breeze *et al.*, 2016). Upon deletion or disruption of the hydrophobic face of this APH region *RTN13* loses the ability to induce constrictions of ER tubules *in vivo* but is still capable of interaction and to form low-mobility oligomers in the ER membrane (Breeze *et al.*, 2016).

However, many interactions of the cytosolic N-terminal domains are still unknown, and it is possible they confer additional functions to the proteins (Tolley *et al.*, 2008). Within the proteins with additional N-terminal domain (RTN17-21) only RTN19 and 20 have N-terminal domains that are predicated to be involved in sterol biosynthesis. For RTN17, 18, and 21 no conserved domain is predicted at the N-terminus (BLASTP).

Reticulons in yeast have been shown to interact with other proteins, for example dynamin-related GTPases (Hu *et al.*, 2009), and the ability of the RTN N-terminal functional domain to facilitate protein-protein interactions may allow the reticulons to tether such molecules to the ER membrane (Nziengui *et al.*, 2007). This could be a potential role for the extended N-terminus of reticulons RTN17, 18 and 21 for which no functions have yet been predicted or identified.

RTN19 features a predicted 3BETAHSD/D domain with a predicted decarboxylatingsterol-4-alpha-carboxylate 3-dehydrogenase activity (BLASTP, AraCyc) indicating a role in sterol biosynthesis. Overexpression of this enzyme, prior to it being identified as a reticulon, in a yeast *erg26* mutant background complemented the mutant (Rahier *et al.*, 2006) which again points towards a role in sterol biosynthesis for RTN19. The closest reticulon homologue to RTN19 is *RTN20* which also features a predicted decarboxylating 3β-hydroxy-4α- methylcholestenecarboxylate 3-dehydrogenase enzymatic domain (AraCyc).

Both these reticulon proteins show high homology in the N-terminal domain to a third protein, 3BETAHSD/D1, with involvement in sterol biosynthesis but without any transmembrane or reticulon homology domain. Like RTN19, 3BETAHSD/D1 is also capable of complementing the yeast *erg26* and in *in vitro* assays with yeast microsomal fractions both enzymes showed 3BETAHSD/D activity with a wide range of steroid substrates (Rahier *et al.*, 2006).

*3betahsd/d1* and *rtn19* single as well as double mutants display no visible phenotype during the plant life cycle (Kim *et al.*, 2012) indicating that other proteins such as *RTN20* or another close homologue At2g33630 may be capable of carrying out the same function in sterol biosynthesis. On the other hand overexpression of 3BETAHSD/D1 and RTN19 appears to affect auxin transporter activity and shows reduced responsiveness to auxin efflux inhibitor NPA (Kim *et al.*, 2012); it is discussed that this could be due to alterations in the sterol composition in the membranes in these overexpression lines (Kim *et al.*, 2012).

Here we show that *rtn20* and *rtn19* mutants display significant changes in the sterol content of arabidopsis roots (Fig. 7), which could likely be due to the functions of the additional N-terminal domain.

### Subcellular localisation/expression pattern

All plant reticulons described so far are preferentially associated with ER tubules and the curved edges of cisternae. Overexpression of reticulons results in severe constrictions of ER tubules and reticulons are able to convert ER membrane sheets into tubules (Tolley *et al.*, 2008, 2010; Sparkes *et al.*, 2010).

RTN19 tagged to a fluorescent protein also labels the ER (Fig. 4) but is not restricted to tubules as shown for other reticulons (Sparkes *et al.*, 2010). RTN20 displays a rather uncharacteristic expression pattern by labelling what appears to be potential ER-membrane subdomains (Fig. 3). RTN19 but not RTN20 features a dilysine (KKXX) ER retrieval motif (Vincent *et al.*, 1998; Benghezal *et al.*, 2000) at the very C-terminus. This motif promotes the retrieval of type I membrane proteins from the Golgi apparatus back to the ER. It was described before though that ER retrieval motifs are non-functional when introduced near an APH region in the cytoplasmic tail (Vincent *et al.*, 1998). For RTN 20 the C-terminal region after the TMDs consists of a significantly higher ratio of hydrophobic amino acids (49%) than for RTN19 which only has 32% of hydrophobic amino acids. This results in a strong hydrophobic face for RTN20 which might explain the special pattern of this protein on the ER membrane. Other than e.g. RTN13 that features a C-terminal amphipathic helix with a hydrophobic face opposite a hydrophilic face (Breeze *et al.*, 2016) such an amphipathic helix cannot be predicted for RTN20.

3BETAHSD/D1 colocalises with the ER exit site markers Sec16 and SAR1A (Fig. 10). 3BETAHSD/D1 does not have any predicted ER signal sequences or TMDs but, similar to some of the reticulon proteins, features an ER retrieval motif. ER exit sites (ERES) are specialized regions of the ER where secretory proteins are concentrated and leave the ER for export to the Golgi bodies (Aridor *et al.*, 1999, 2001; Hammond and Glick, 2000). ERES are characterized by local accumulations of COPII proteins such as Sec16, together with the dimers Sec23/24 and Sec31/13 (Budnik and Stephens, 2009; Marti *et al.*, 2010; Miller and Barlowe, 2010). The localisation of 3BETAHSD/D1 on ERES is consistent with the finding of enzymatic activity for this enzyme in yeast microsomal fractions but not cytosolic fractions (Rahier *et al.*, 2006) although the targeting to the ER is unclear as 3BETAHSD/D1 is lacking the typical signal sequences or TMDs. Proteins homologous to 3BETAHSD/D1 were identified in yeast and mammals, both of which were ER localised and have roles in the sterol biosynthetic pathway (Gachotte *et al.*, 1998; Caldas and Herman, 2003). The lack of predicted transmembrane regions within these proteins suggested they are only peripherally associated with the ER and instead tethered via interactions with other ER-associated proteins (Gachotte *et al.*, 1998). This could also be the case for 3BETAHSD/D1.

### Lipid composition/function in sterol biosynthesis

Phytosterols play major roles in plants not only as structural membrane molecules (Hartmann, 1998) but also for plant growth and development (Clouse, 2002) and hormone signalling (Lindsey *et al.*, 2003; Men *et al.*, 2008; Mongrand *et al.*, 2010; Simon-Plas *et al.*, 2011). For the biological function of these sterols the removal of the two C-4 methyl groups is crucial. It has been hypothesised that ER tubules are sites for lipid production (Friedman and Voeltz, 2011), and our mutant data indicates that RTN19 and 20 could well be involved (Fig. 7).

Considering the various subcellular localisations of the three potential enzymes, there is the possibility that they also have different functions. It has been shown in yeast that Erg28p can work as an ER transmembrane scaffold protein that tethers several partners of the C-4 demethylation enzymatic complex (Mo *et al.*, 2002; Mo and Bard, 2005). In addition, it was found that Erg28 in yeast was more closely associated with some Erg proteins than others, indicating the possibility of different levels/types of interactions between these proteins (Mo and Bard, 2005). Such different levels of interactions could then correspond to different roles in regulation of metabolism, and in different ER domains for example involved in the formation of different structures or membrane interactions (ERES, lipid and protein bodies, membrane contact sites between ER and plasma membrane, and mitochondria etc.). Therefore, we could consider different roles of sterol metabolism in these different mechanisms.

In arabidopsis AtErg28 tethers the sterol C-4 demethylation complex to prevent the accumulation of sterol intermediates that can interfere with the regulation of the polarized transport of auxin (Mialoundama *et al.*, 2013). Since such a link between the biosynthesis and/or the regulation of the biosynthesis of sterols was established to have an impact on a specific function (for example auxin transport), the following possibilities for RTN19, RTN20 and 3BETAHSD/D1 could be considered: 1) The proteins interact with different enzymatic complexes and are engaged in different enzymatic activities. 2) They work as different ER transmembrane scaffold proteins to tether different partners and therefore regulate different metabolic pathways as AtErg28 does. 3) They can also interact with other types of proteins to regulate the formation of different structures/domains of various membranes where sterol biophysical properties are required.

We could speculate for example whether RTN19 works more on bulk sterol biosynthesis in the ER network and therefore might be involved in any enzyme activity whereas RTN20 and 3BETAHSD/D1 proteins are more specifically focused on the regulation of lipid/sterol biosynthesis linked to ERES formation and/or COPII vesicle fission in ER domains. At the same time each of the proteins could compensate for the others. Therefore, to unravel the exact roles of these proteins, the putative enzyme activities and/or the search of the putative partners will have to be investigated.

## Methods

### Bioinformatics analysis

The functional domains of the reticulon family of proteins from Arabidopsis thaliana were analysed. The protein ATG numbers were searched for in TAIR and a protein BLAST in the *A. thaliana* database done to ensure all 21 sequences were correct and any splice variants of the proteins identified. A total of 35 sequences were obtained and phylogenetic analysis was performed with a one-click analysis using phylogeny.fr (Dereeper *et al.*, 2008; Dereeper *et al.*, 2010). For phylogenetic analysis protein BLAST of RTN19 and 20 was performed within the yeast and human databases to identify homologous proteins. The sequences obtained were then used to search for additional arabidopsis proteins by doing a protein BLAST in the *A. thaliana* database. Phylogenetic analysis of the identified sequences was done using a phylogeny.fr one-click analysis, with the RTN17, 18 and 21 proteins used as outgroup.

Membrane topology with hydrophobic membrane-spanning regions was analysed using TOPCONS (Bernsel *et al.*, 2009).

### Cloning of expression plasmids

Primers were obtained from Eurofins Genomics. Q5 high-fidelity DNA polymerase (New England Biolabs) was used for all polymerase chain reaction reactions. Genes of interest were cloned into the modified binary vector pB7WGF2 containing an N-terminal or pB7FWG containing a C-terminal green fluorescent protein (GFP) and pB7RWG2 or pB7WGR2 for the red fluorescent protein (RFP) (Karimi *et al.*, 2005) using Gateway technology (Invitrogen).

### Plant material and transient expression in tobacco leaf epidermal cells

For *Agrobacterium*-mediated transient expression, 5-week-old tobacco (Nicotiana tabacum SR1 cv Petit Havana) plants grown in the greenhouse were used. Transient expression was carried out according to Sparkes *et al.* (2006).

In brief, each expression vector was introduced into the *Agrobacterium* strain GV3101 by heat shock transformation. Transformant colonies were inoculated into 5 ml of YEB medium (5 g/l beef extract, 1 g/l yeast extract, 5 g/l sucrose and 0.5 g/l MgSO_4_. 7H_2_O) with 50 μg/ml spectinomycin and rifampicin to keep the selection pressure up. The bacterial culture was incubated overnight in a shaker at 180rpm 25°C. 1 ml of the bacterial culture was pelleted by centrifugation at 2200 *g* for 5 min at room temperature. The resulting pellet was washed twice with 1 ml of infiltration buffer (50 mM MES, 2 mM Na_3_PO_4_.12H_2_O, 0.1 mM acetosyringone and 5 mg/ml glucose) and then resuspended in 1 ml of infiltration buffer. The bacterial suspension was diluted to a final OD_600_ of 0.1 and carefully pressed through the stomata on the lower epidermal surface using a 1 ml syringe.

Transformed plants then were incubated in a growth cabinet at 22°C for 48 h. Samples from transient expression and transformed arabidopsis plants (cotyledons) were imaged using a 100x/1.46 NA oil immersion objective on a Zeiss LSM880 equipped with an Airyscan detector. For imaging of the GFP–RFP combinations, samples were excited using 488 and 543 nm laser lines in multi-track mode with line switching. Images were edited using the ZEN image browser.

### FRET-FLIM Data Acquisition

Tobacco Epidermal leaf samples of infiltrated tobacco plants were excised, and FRET-FLIM data capture was performed according to Schoberer and Botchway (2014) using a two-photon microscope at the Central Laser Facility of the Rutherford Appleton Laboratory. GFP and mRFP expression levels in the plant samples within the region of interest were confirmed using a Nikon EC2 confocal microscope with excitation at 488 and 543 nm, respectively. A 633-nm interference filter was used to minimize the contaminating effect of chlorophyll autofluorescence emission. A two-photon microscope built around a Nikon TE2000-U inverted microscope was used with a modified Nikon EC2 confocal scanning system to allow for multiphoton FLIM (Botchway *et al.*, 2015). 920 nm laser light was produced by a mode-locked titanium sapphire laser (Mira; Coherent Lasers), producing 200-fs pulses at 76 MHz, pumped by a solid-state continuous wave 532-nm laser (Verdi V18; Coherent Laser). The laser beam was focused to a diffraction limited spot through a water-immersion objective (Nikon x60 VC; 360, numerical aperture of 1.2) to illuminate the specimen. Fluorescence emission was collected without descanning, bypassing the scanning system, and passed through a BG39 (Comar) filter to block the near-infrared laser light. Line, frame, and pixel clock signals were generated and synchronized using a fast microchannel plate photomultiplier tube as external detector (Hamamatsu R3809U). Linking these via a time-correlated single-photon-counting PC module SPC830 (Becker and Hickl) generated the raw FLIM data. Data were analysed by obtaining excited-state lifetime values of a region of interest. Calculations were made using SPC Image analysis software version 5.1 (Becker and Hickl). The distribution of lifetime values within the region of interest was generated and displayed as a curve. Only values with a X^2^ between 0.9 and 1.2 were taken into account. The median lifetime value in the region of interest was taken to generate the range of lifetimes per sample. At least three nuclei from a minimum of three independent biological samples per protein-protein combination were analysed, and the average of the ranges was taken.

### Reverse transcriptase-PCR

RNA was isolated using TRIzol® (Thermo Fisher Scientific) according to the manufacturer’s instructions. The AMV First Strand cDNA Synthesis Kit (New England Biolabs) was used according to the manufacturer’s instructions to create cDNA for wildtype arabidopsis as well as for *rtn20* and *rtn6* mutants. The resulting cDNA was probed with primers for full-length products of RTN20 and RTN6, respectively using Q5® High-Fidelity DNA Polymerase (New England Biolabs).

### Lipid analysis

Lipids from roots and leaves of two week-old seedlings (wild type and mutant lines) were extracted by grinding the tissues in a mixture of chloroform/methanol (2:1, v/v) at room temperature. Then lipid extracts were washed two times with 9 ‰ NaCl (1/4 of the organic solvent volume). The organic solvent was then evaporated and lipid extracts were dissolved in an appropriate volume of chloroform/methanol (1:1, v/v). Phospholipids were analyzed by loading total lipids onto HPTLC plates (60F254, Merck, Darmstadt, Germany), which were developed in methyl acetate/n-propanol/chloroform/methanol/0.25% aqueous KCl (25:25:25:10:9, v/v) according to Heape et al. (1985). To isolate and quantify sterols, total lipids were loaded onto HPTLC plates developed with hexane/ethylether/acetic acid (90:15:2, v/v) as in Laloi et al (2007). Lipids were identified by co-migration with known standards and quantified by densitometry analysis (Macala *et al.*, 1983) using a TLC scanner 3 (CAMAG, Muttenz, Switzerland) as described in Laloi et al. (2007).

### EM fixation and data acquisition

Arabidopsis root segments were fixed in 1% glutaraldehyde and 1% paraformaldehyde in 0.1M sodium cacodylate buffer (pH 6.9) for 60 min, washed in buffer and poststained for 12hr in a mixture of zinc iodide and 1% osmium tetroxide (Hawes *et al.*, 1981), followed by ethanol dehydration and embedding in Spurr resin (hard). Resin blocks were mounted onto 3View stubs with conductive epoxy glue (Chemtronics) and left to harden overnight. Blocks were trimmed and sections with a Gatan 3View system and SBFSEM images were collected with a Zeiss Merlin Compact field emission SEM. Slice thickness was 50-70 nm and the block face imaged under variable pressure (20-55 pa) at 3-4 keV with a pixel dwell time of 3 μs and pixel size of 4-6.2 μm. Initial image handling and alignment was carried out using *etomo* (IMOD, Boulder, Colorado (Kremer *et al.*, 1996). Segmentation and reconstructions was achieved using Amira (Version 6.2, FEI, Eindhoven).

### Accession numbers

Sequence data for the genes mentioned in this article can be found in the GenBank/EMBL databases using the following accession numbers:

AT4G23630 (RTN1); AT4G11220 (RTN2); AT1G64090 (RTN3); At3g61560 (RTN6); At2g20590 (RTN17); At4g28430 (RTN18); At2g26260 (RTN19, 3BETAHSD/D2); At2g43420 (RTN20); At5g58000 (RTN21); AT1G47290 (3BETAHSD/D1).

## Acknowledgements

This research was supported by a 13 ERA-CAPs grant to CH (BBSRC BB/M000168/1) and BBSRC ALERT13 funding (BB/C014122/1). The authors thank Guillaume Bouyssou for excellent technical assistance with the lipid analysis.

## Author contributions

VK and CH designed the project. LMP and PM carried out the lipid analysis. JU and VK performed the bioinformatics analysis. VK cloned the constructs, carried out confocal microscopy and FRET-FLIM, and created the plants. SWB and VK analysed the FRET-FLIM data. LH, JR, MK and CH carried out the tissue preparation for electron microscopy. LH acquired and analysed the electron microscopy data. VK, PM and CH wrote the manuscript.

